# Cell cycle criticality as a mechanism for robust cell population control

**DOI:** 10.1101/2025.06.12.659375

**Authors:** Benjamin D. Simons, Omer Karin

## Abstract

Tissue homeostasis requires a precise balance between cellular self-renewal and differentiation. While fate decisions are known to be closely linked to cell cycle progression, the functional significance of this relationship is unclear. Here, we develop a mechanistic framework to analyse cellular dynamics when cell fate is coupled to cell cycle length. We focus on a distinct feature of cell cycle regulation where mitogens act as control parameters for a bifurcation governing the G1-S transition. Under competitive feedback from cell-cell interactions, the cell cycle regulatory network fine-tunes to the critical point of this bifurcation, becoming highly sensitive to mitogenic signalling. This critical positioning lengthens G1 while amplifying cell-to-cell variability in signalling and biochemical states. Such regulation confers significant advantages for controlling cell population dynamics, including maintaining a robust population set-point and rejecting mis-sensing mutants. The mutant rejection capability trades off against tissue growth and repair. Counter-intuitively, we propose that adult stem cells couple prolonged G1 with increased self-renewal propensity to efficiently eliminate mis-sensing mutants. Our theory explains and predicts regulatory patterns across development, homeostasis, and ageing.

## Introduction

Tissues are formed and maintained through a dynamic interplay of cell division, differentiation, and loss. Achieving the correct tissue size and composition requires a precise balance of these processes. Many tissues, such as blood and renewing epithelia, are sustained by multipotent stem and progenitor cells [1–3]. In such tissues, cell loss or changes in the demand for specific cell types trigger compensatory processes of cell division and differentiation within the stem and progenitor populations, thereby restoring tissue homeostasis. Similarly, in developing tissues, progenitor populations must modulate their division and differentiation rates as the tissue grows to attain the appropriate size and composition. The problem of cell population control is also crucial for engineering synthetic circuits with applications in industry and biomedicine [4–7].

Functionally, cell fate control circuits must rapidly and precisely correct deviations from set points while suppressing the expansion of mutants that mis-sense signals related to cell division or removal. Experimental and theoretical work has demonstrated that cell-cell negative feedback plays a central role in this regulation [2, 8, 9]. In this mechanism, population composition modulates the local microenvironment through biochemical or mechanical feedback, thereby influencing subsequent fate decisions through the modulation of intracellular regulation mechanisms. However, the design principles of the intracellular mechanisms that ensure the robust and efficient operation of these population control circuits remain unclear.

In this study, we propose that cells employ a *temporal* mechanism for cell fate regulation, where cells exploit a coupling between the cell division rate and cell fate choice to implement robust cell population control. While the processes regulating these rates may, in principle, be independent, in many systems they appear to be tightly coupled. The regulation of the cell division rate occurs primarily through controlling the average length of the G1 phase until transition into the S phase, in a process known as cell cycle entry. Major signalling pathways that control cell division rate through adjustment of the duration of the G1 phase of the cell cycle (mitogens), such as Wnt/β-catenin and MAPK/ERK, are also key regulators of cell identity [10–15]. Increasing experimental evidence suggests that the length of G1 itself controls cell identity and differentiation [16–19]. This regulatory mechanism has been demonstrated in various contexts including embryonic stem cells, where G1 length determines fate choice [18, 20–25]; during neurogenesis [26–32]; in adult stem cell populations [33–39]; and in terminal differentiation [19, 40, 41]. Despite its apparent fundamental importance in modulating cell fate choice in tissues, the implications of the coupling between G1 lengthening and cell fate choices as a mechanism for cell population control remain unclear.

Here, we develop a mechanistic framework to analyse this coupling. We model G1 lengthening as a noisy saddle-node bifurcation, where regulatory signals tune a bifurcation parameter to modulate G1 duration. We show that when G1 length is coupled to cell fate by negative feedback, the dynamics leads generically to self-tuning to the vicinity of a critical bifurcation point. This self-tuning results in convergence to a robust fixed point of the population size. The critical mechanism provides the cellular population with rapid responses to deviations from this fixed point and efficient rejection of mis-sensing mutants, explaining experimental observations on cell cycle dynamics. We demonstrate that these two features provide a trade-off with each other and are favoured by alternative topologies, explaining patterns of regulation in development, homeostasis, and ageing.

## Results

### G1 lengthening occurs by critical dynamics that amplifies variation between cells

The G1-S transition is governed by a molecular regulatory network that involves interactions between the proteins Rb, E2F, CycD-CDK4/CDK6 (abbreviated CycD), and CycE-CDK2 (abbreviated CycE) [12, 42–46] (Figure 1A). The transition occurs following an increase in the activity of E2F and the transcription of CycE proteins. The underlying dynamics correspond to a bistable activity pattern due to positive feedback [43]. Active Rb protein forms a complex with E2F that represses its activity and prevents CycE transcription, while CycE in turn, reduces the activity of Rb by phosphorylation, closing the positive feedback loop. The feedback between these proteins can stabilize both a low E2F state, and a high E2F state that is associated with the G1-S transition.

**Figure 1.**
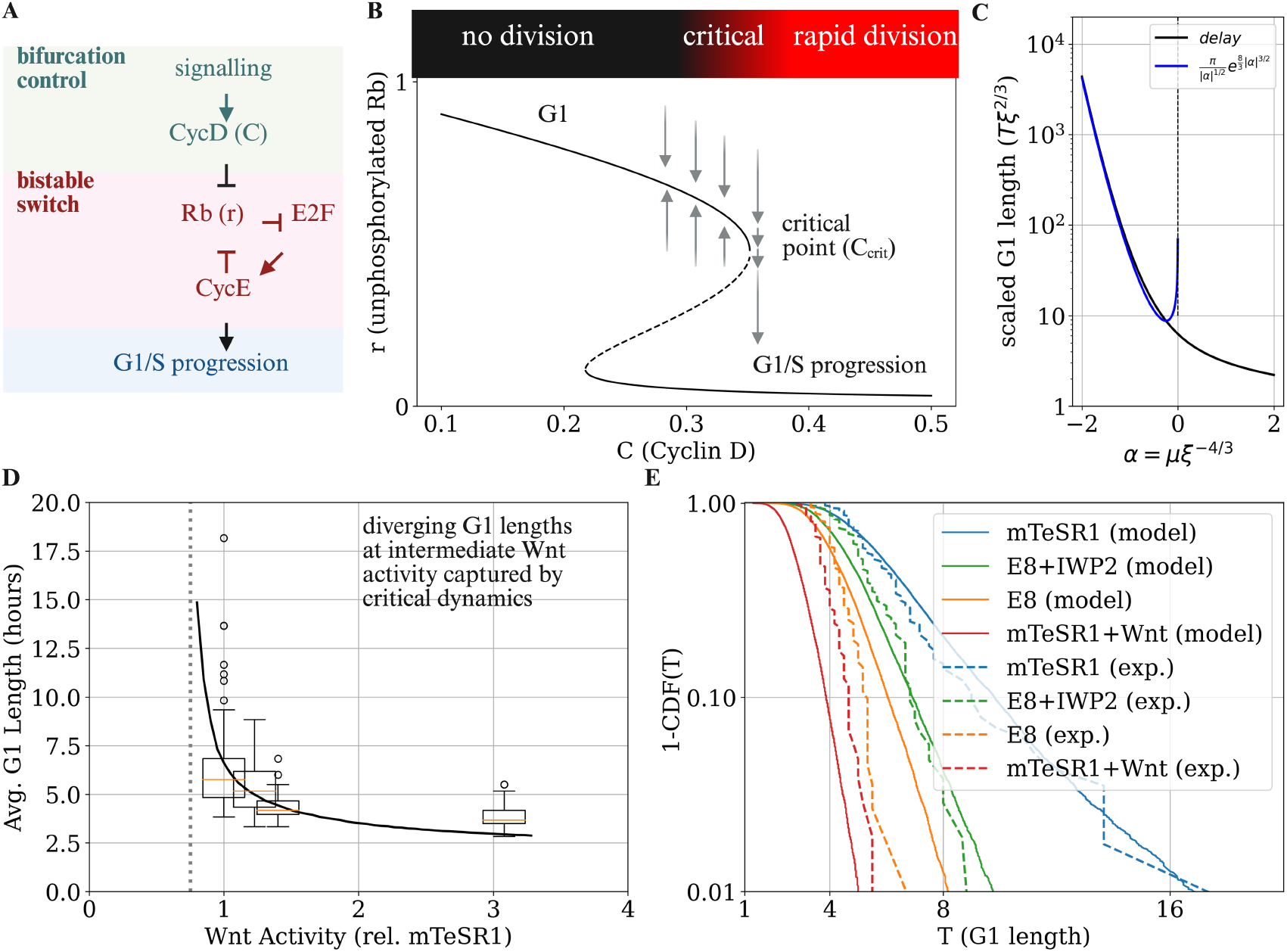
Model for G1 lengthening. (A) Outline of the circuit that controls G1-S progression during cell cycle, with signalling input controlling a positive-feedback based mechanism. (B) Bifurcation diagram for the G1-S progression mechanism showing how changes in CycD concentration mediate an abrupt change of unphosphorylated Rb concentration (see main text). (C) Dependence of the G1 length *T* on the scaled distance from the bifurcation *µ* (black line). For *µ <* 0, this dependence can be captured approximately by a stretched exponential dependence (blue line). (D) G1 length distribution as a function of intracellular Wnt activity (as reported by AXIN2 versus GAPDH expression, relative to lowest reported activity level) for hESCs grown in the following experimental conditions: mTeSR1 media, E8 media supplemented with IWP2 (where IWP2 is a Wnt inhibitor), mTeSR1 media supplemented with Wnt, and E8 media (see main text). The G1 length shows a sharp and highly nonlinear increase associated with increased variability around a specific Wnt activity level (data from Ref. [23]). This sharp increase is consistent with a G1 elongation mechanism based on a noisy saddle-node bifurcation. The black line corresponds to (average) model simulations with the bifurcation control parameter proportional to Wnt activity, taking a critical concentration at 3*/*4 of the minimal observed Wnt activity level (dashed gray line) with fixed cell-to-cell variability in the control parameter (Methods). (E) Comparison of variability in G1 lengths between data from Ref. [23] and model simulations. The model captures the sharp increase in variability associated with G1 lengthening, which is due to critical dynamics amplifying cell-to-cell variation.

The transition from the low E2F state to the high E2F state is also controlled by the activity of another protein, CycD [42–44, 47]. CycD affects the transition of E2F from off to on by also reducing the activity of Rb via phosphorylation [48]. When CycD is high, a bifurcation occurs that destabilizes the low E2F state and transitions the system to the high E2F state, promoting the G1-S transition. CycD levels are nearly constant throughout G1 and are controlled by a wide range of external signals, including through the MAPK and Wnt/*β*catenin pathways [49, 50]. Thus, a low level of CycD extends G1 as it prevents the G1-S transition, while a high level shortens G1.

While the bistable nature of the Rb/E2F switch is well established [43, 51], there is debate regarding the precise biological mechanisms underlying its implementation [50, 52–54]. We therefore chose to focus on capturing the essential dynamics through a minimal framework, noting that our subsequent analysis applies to a much broader class of possible models. Denoting the level of active (unphosphorylated) Rb protein as *r*, its dynamics are governed by a constant production rate and removal through both degradation, at rate *γ*_*D*_, and phosphorylation, at rate *γ*_*P*_. The phosphorylation rate depends on both CycD activity *C* and a feedback loop driven by the level of *r* itself, which we capture phenomenologically by the relation 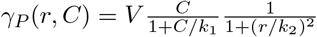 where *k*_1_ controls the scale at which CycD activity becomes saturated, *k*_2_ fixes the level at which Rb modulates its phosphorylation rate, and *V* fixes the net amplitude. We assume that the feedback of *r* on itself, which occurs through the E2F/CycE cascade, is cooperative. We also incorporate the impact of noise, such as that arising from stochastic fluctuations in the levels of different factors that control Rb phosphorylation. These fluctuations can be modelled as a Wiener-type process with a multiplicative noise amplitude *σr* (that we subsequently generalize). The dynamics of *r* are therefore given by the stochastic differential equation,

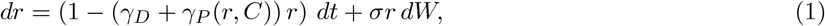

where *dW* denotes a Wiener process (Figure 1B).

Within this framework, the essential dynamics of the circuit are captured by a saddle-node bifurcation, with the activity level of CycD, *C*, serving as a control parameter for the bifurcation. The bifurcation, which is depicted graphically in Figure 1B, is the key component of a wide range of mathematical models for the G1-S transition, from simple one-dimensional models to highly elaborate mechanistic models [43, 45, 55, 56] (Methods). The control parameter *C* (affected by mitogenic signals) destabilizes the state associated with G1, resulting in the G1-S transition. When levels of *C* are low, the system is positioned in a stable configuration (G1) and is resistant to fluctuations. At higher levels of *C*, this phase becomes metastable with the emergence of a second fixed point of the dynamics. Eventually, at a critical level of CycD activity *C* = *C*_crit_, the system transitions rapidly into S phase. Importantly, we assume that *C* is generally constant during G1, and later consider the case where *C* changes in a time-dependent manner.

The saddle-node bifurcation has three key properties that will play an important role in our analysis. The first is that in the vicinity of *C* ≈ *C*_crit_, the dynamics of the system are captured by a one-dimensional noisy process,

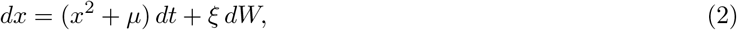

where *x* denotes the rescaled coordinates that parametrize the bistable switch, the time *t* is measured in rescaled coordinates, and *µ* ∝ *C* − *C*_crit_ is the (rescaled) distance from bifurcation. In the context of cell cycle regulation, G1 corresponds to negative *x*, while the G1-S transition occurs at positive *x*. The noise amplitude *ξ* corresponds to the fluctuations near the critical point. The length of G1, denoted *T*, is captured by the typical time taken for *x* to change sign from negative to positive.

The second property is the strong sensitivity of the length of G1, *T*, around the critical point, which is marked by an abrupt, super-exponential increase at the critical threshold (with log *T* ∼ |*µ*|^3*/*2^) [57–59] (Figure 1C). As such, any prolonged G1 length is predicted to occur when the regulatory network governing the G1-S transition is poised in the vicinity of the critical point *C* ≈ *C*_crit_, with *µ* ≈ 0, where the dynamics are captured by Eq. (2). The third property relates to the variation between individual cells and between signalling and media conditions. Different media conditions are associated with different levels of mitogenic signalling. Cellular heterogeneity introduces additional variation arising from differences in cellular physiology, biochemical composition, and spatial configuration. These variations are captured through variation in the coordinate *µ*. Crucially, the steep lengthening of *T* and its strong sensitivity to *µ* around the critical point (Figure 1C) amplifies cell-cell variation and variation between conditions, resulting in steep and divergent G1 lengthening around *µ* ≈ 0.

To test whether experimental data is consistent with G1 lengthening occurring through a noisy saddle-node bifurcation, we analysed published data on the distribution of G1 duration and its regulation in human embryonic stem cells (hESCs) [23] (Figure 1D,E). G1 phase length in hESCs depends on culture media conditions, including signalling factor concentrations, such as Wnt signalling through the Wnt/β-catenin pathway [18, 20–24].

In our modelling framework, activation of the Wnt pathway affects the control parameter *C* (Figure 1D). The model therefore predicts a steep and divergent lengthening of G1 around a specific critical level of Wnt activity. Indeed, these predictions are consistent with the experimental findings of Jang et al. [23]. After culturing hESCs in media conditions associated with varying degrees of Wnt activation, they observed that the decrease in average G1 length occurs sharply within a narrow range of endogenous Wnt activity (Figure 1D), with a functional dependence consistent with a bifurcation event [23].

The model also explains observations on cell-to-cell variation under different growth conditions (Figure 1E). In E8 media, hESCs exhibit a G1 length distribution tightly clustered around an average of *T* ≈ 4.3 hours. In contrast, in mTeSR1 media, the average G1 duration increases to *T* ≈ 6.3 hours. This approximately 50% increase in mean *T* is accompanied by a much larger increase in variability (translating to an approximately 250% increase in the coefficient of variation) due to the emergence of a heavy tail of cells with very long G1 duration in mTeSR1 media, associated with G1 lengths exceeding 6 hours. Similar effects were observed when G1 length was perturbed by treatment with Wnt or Wnt inhibitors. The model captures this dramatic change in variability, where lengthening is caused by critical dynamics that amplify variation. Collectively, these findings support a model in which G1 length is extended by a (noisy) saddle-node bifurcation.

### Tuning of G1 length by competition provides a set-point for self-renewing tissues

The observation that cell fate decisions may require specific lengthening of G1 raises questions about how this lengthening is achieved in tissues. We focus here on a scenario in which a cell, such as a tissue stem cell, must select between self-renewal and differentiation through symmetric cell divisions, with this decision depending on the duration of G1 length *T*. (Note that the model can readily generalize to other patterns of cell fate, such as choosing between alternative fates in developmental settings.) The strong sensitivity of G1 length on the bifurcation parameter *C*, and consequently to external signals such as Wnt signalling, suggests that for an arbitrary parametrization, G1 length will typically be either very short or effectively infinite, implying a strong imbalance between self-renewal and differentiation. Denoting by *N* the number of undifferentiated cells in the population, we can capture the net growth of *N* by:

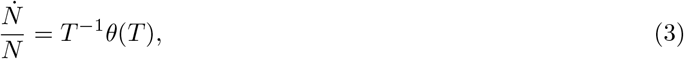

where *θ* captures the dependence of cell fate on the cell cycle.

To see how G1 lengthening can occur, we note that cells typically share and modulate environmental signals, including the concentrations of the molecules that adjust G1. This results in competition and negative feedback, where a change in the abundance or composition of the cell population by G1 lengthening or shortening feeds back on the environmental signal. As an example, consider the case where the cell population size or density *N* modulates the control parameter for the bifurcation,

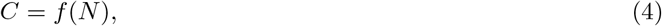

where *f* may correspond to the effect of competition over secreted factors and may be affected by contact inhibition. For simplicity, here we assume that differentiated cells exit the population (though this assumption is not essential, as the dynamical patterns we discuss also emerge in more complex regulatory circuits). In this case, a balanced state can occur when *N* is constant,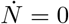. Here, the G1 duration *T* = *T*_*S*_, where the growth rate of *N* is zero, *θ*(*T*_*S*_) = 0. Since every finite *T*_*S*_ occurs at the vicinity of *C* ≈ *C*_crit_, where the cell cycle length is prolonged, the set-point of the self-renewing population becomes

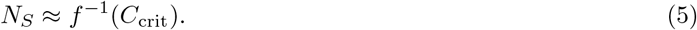

This fixed point is stable if around the fixed point we have that

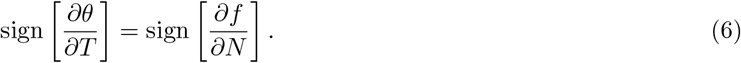

This equality holds for two distinct topologies. In the first case, an increase in cell abundance prolongs G1, requiring that self-renewal preferentially occurs at shorter G1 lengths. This topology will henceforth be referred to as the **Renewal precedes Differentiation** (RD) topology (Figure 2A). The RD topology could be implemented in cells, for example, through a slow G1-dependent accumulation of a differentiation factor, as occurs in adipocytes [19], or through G1-dependent susceptibility to receiving differentiation signals. Alternatively, an increase in *N* may preferentially *shorten* the length of G1, coupled with increased differentiation at shorter G1 lengths. This can occur if cells need sufficient time in G1 to maintain their identity, such that early entry into the cell cycle increases the likelihood of differentiation. For example, if a differentiating factor is degraded in G1 and accumulates in other cell cycle phases, or through G1-dependent susceptibility to signals required for maintenance of cell identity. We denote the second case as the **Differentiation precedes Renewal** (DR) topology (Figure 2B).

**Figure 2.**
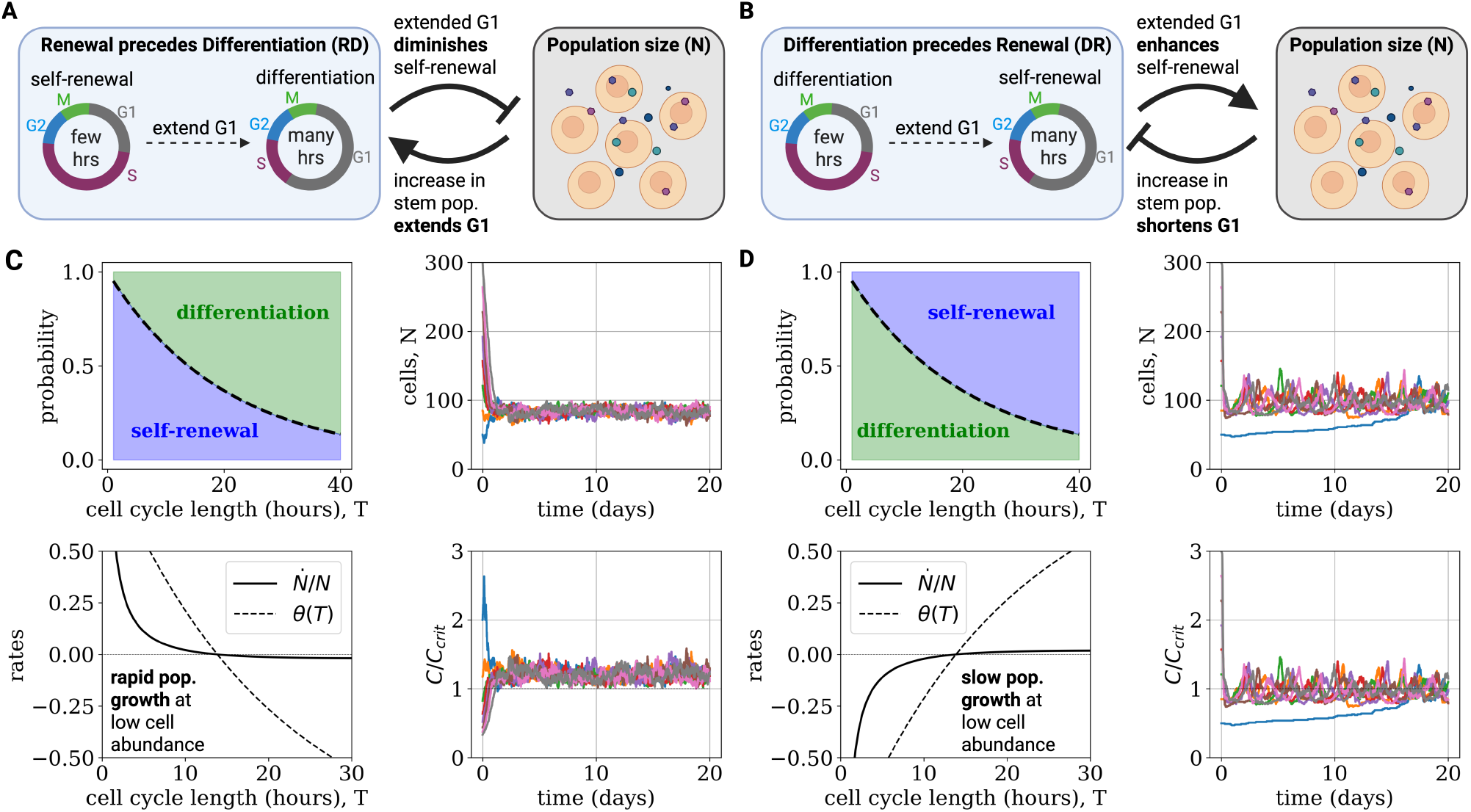
Set point regulation and alternative topologies. (A) RD topology, with rapid cell division favouring self renewal. (B) DR topology, with rapid cell division favouring differentiation.(C) Simulations of population dynamics under the RD topology from different initial concentrations, with feedback set by *C* = *f* (*N*) = *C*_crit_*N*_st_*/N* with *N*_st_ = 100, demonstrating convergence to the critical set point (right subpanels). In the left subpanels, we plotted the probability that a cell division will result in duplication (self-renewal) versus differentiation as a function of cell cycle length (*T*) and the effective rate functions *θ*(*T*) and 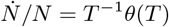 (see main text). To account for additional processes that can drive cell division, such as injuries or physiological changes, all simulations included, in addition to regulated cell divisions, random fixed-rate cell divisions, which were set at a rate of *λ* = 1*/*30 days. (D) Corresponding circuit and simulations for the DR topology. Note the slow recovery at low initial population, which is driven by random cell divisions.

While both the RD and DR topologies exhibit sensitive responses to changes in cell abundance and signalling factors, they have very different dynamics away from the set point. Consider the case where the number of cells *N* is much lower than the set point. For both topologies, this can result in pure self-renewal, but the dynamics differ. In the RD topology, self-renewal occurs rapidly due to the short cell cycle time (fixed effectively by *T*). Only as the cell population approaches its steady state does a slowdown in G1 occur, resulting in large-scale cell cycle arrest (Figure 2C). In control theory, this regulation is known as *bang-bang control*, providing the optimal response for achieving the cell population set point in minimal time [60]. The dynamics predicted by the RD topology appear to be widespread in developmental settings [26, 60–62]. The DR topology, on other hand, exhibits sluggish responses when cell abundance is low due to the extension of cell division times (Figure 2D). We, thus, propose that the RD topology may provide an optimal mechanism for rapid tissue development.

### Mutant rejection versus growth regulation determines design principles of tissue circuits

Within a steady-state environment, the fate decisions of wild-type cells are such that cell divisions are equally likely to result in differentiation or self-renewal. In such an environment, mutant cells that mis-sense input signals in such a way that they perceive the population size as too low would be biased towards self-renewal. This bias leads to their expansion within the population, ultimately displacing the wild-type cells and shifting the system’s set-point (Figure 3A). Preventing such mutant takeover is therefore a fundamental challenge for population-control circuits.

**Figure 3.**
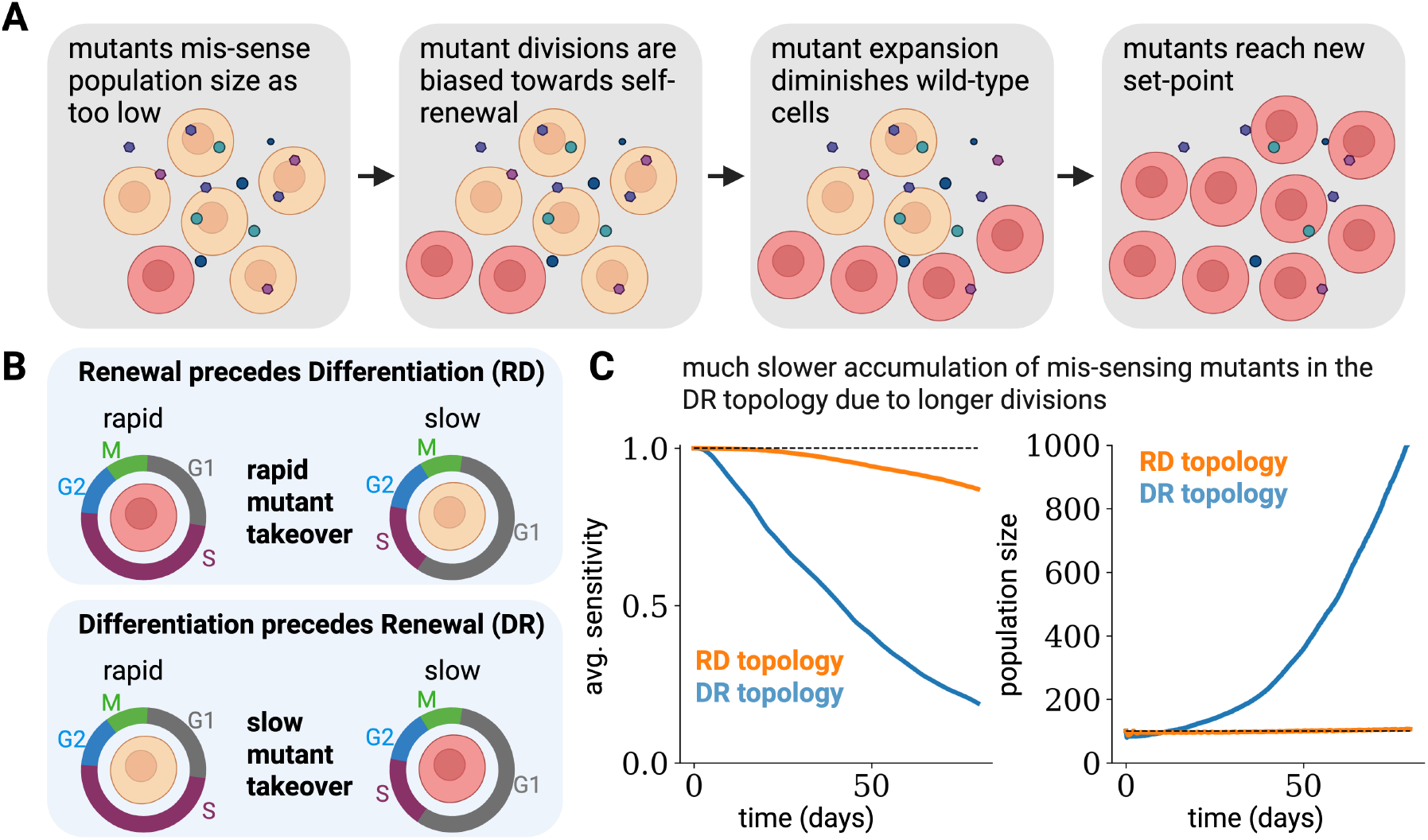
Efficient mutant rejection in the DR topology. (A,B) RD and DR dynamics show differential dynamics, with the sharp sensitivity around the critical point ensuring that mis-sensing mutants will either have rapid or prolonged cycling times, respectively. (C) Population dynamics were simulated with mutations as follows. *f* (*N*) was replaced in the simulations by *f* (*α*_*i*_*S*), with *α*_*i*_ denoting the sensitivity of cell *i*, initialized at unity. At each cell division, *α*_*i*_ was “mutated” with probability *p*_*m*_ or inherited with probability 1 − *p*_*m*_, with mutation corresponding to multiplication of *α*_*i*_ by a random number drawn from a distribution ℱ. (Here, ℱ was taken as a log-normal distribution whose log-average is zero and log-standard derivation is taken as 0.5, and *p*_*m*_ = 0.001.) Both the RD and DR topologies drift towards mutants that under-sense the population size and thus have a higher population fixed point. However, the DR topology is more resistant, as its invading mutants are associated with longer cell cycle durations. We set the minimal sensitivity to *α*_*i*_ = 0.03.

Formally, within the modelling framework, mutants can have altered sensing or processing of growth signals, leading to an altered mapping of cellular composition to growth regulation, captured by a dependence of the control parameter on *N*,

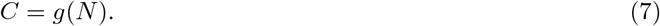

This results in a different population set-point for the mutants *N* ^*′*^ = *g*^−1^(*C*_crit_).

The strong sensitivity of cell fate around the critical point provides an additional benefit: the efficient rejection of “cheater” mutants that have aberrant sensing or processing of regulating signals. To see how the sensitivity in the critical region contributes to mutant rejection, consider the case where an individual mutant is induced on the steady-state background of wild-type cells *N* = *N*_*s*_. These mutants will have either a very short G1 (*T* on the order of a few hours) or a very long G1 length. Efficient mutant rejection only requires that the growth rate *T* ^−1^*θ*(*T*) will be low at short G1 and long G1 lengths. As we will see, this requirement manifests differently for the DR and RD topologies.

The condition for efficient mutant rejection is achieved naturally in the DR topology (Figure 3B,C). Here, mutants with shorter G1 length are lost due to differentiation. Mutants with an extended G1 phase may accumulate, but this process occurs over a long timescale due to their much slower division rate. Consequently, the model predicts a gradual accumulation of clones with impaired responsiveness to signals that typically shorten the G1 phase duration for this regulatory topology.

In contrast, the RD topology is fragile to the invasion of mutants (Figure 3B,C). Mutants that are associated with a shorter G1 phase have both a shorter cell cycle (and thus, most likely, a faster overall division rate) and a higher self-renewal potential. As these mutants expand in cell number, they may alter their environment in such a way that they confer a growth disadvantage on their neighbouring wild-type cells, eventually resulting in the elimination of the wild-type population.

There are several ways that evolved systems may mitigate the detrimental aspects of mutant clone invasion associated with the RD topology. One possibility is that there is an overall compensation of S, G2, and M lengths to shorter G1, which indeed appears to be the case for embryonic stem cells [23]. Another possibility is to engineer biphasic regulation of the growth by G1 length, where very short G1 lengths can result in cell removal (Figure 3C), due, for example, to insufficient time to prepare for DNA replication [63]. In this case, mutants with very short G1 will be eliminated. Biphasic regulation leads to the emergence of an additional unstable fixed point at short G1 lengths, and can thus result in a runaway elimination of the cell population following large perturbations [8], which can be detrimental for tissue development or regeneration.

The advantages of the DR topology for mutant rejection suggest that it may offer significant benefits for regulating self-renewal in tissues requiring long-term replenishment. It is well-established that in many such tissues, including the human blood, skin, and gut, self-renewal depends on a slow-cycling stem cell population with long-term self-renewal potential. This population transitions into a more rapid-cycling transit-amplifying (TA) population with limited self-renewal potential [64]. For the DR topology to play a role in the dynamics of these systems, one would expect that a shortening of the G1 length would *drive* entry from the stem to the TA compartment. While not conclusive, evidence suggests that, across several self-renewing tissues, genetic changes that increase proliferation serve to diminish the size of the stem cell pool, whereas inhibition of proliferation enhances self-renewal at the expense of cellular differentiation [33–35, 65–67]. Similarly, during the ageing of self-renewing tissues, there appears to be an accumulation of cells with improved self-renewal coupled to diminished cell division rate [68–70].

The DR topology is predicted to be associated with the accumulation of slow-cycling mutants with high capacity for self-renewal (Figure 3C). One system where clonal dynamics have been characterized during ageing is the blood, where a few clones with specific mutations accumulate during ageing, in a phenomenon known as clonal haematopoiesis [71]. The model specifically predicts that this process will be driven by cells that have prolonged cell cycle duration and that are associated with a higher set point 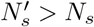. Both predictions appear to hold for the haematopoietic system as, with age, haematopoietic stem cells appear to grow in number [72, 73], have longer cell cycle times driven by a longer G1 phase [70], and are less sensitive to mitogens [70]. This suggests the possibility that ageing of haematopoietic stem cells is driven by competition and the accumulation of slowly dividing cells with impaired sensitivity, which are hallmarks of the DR topology.

The RD topology, on the other hand, appears to be widespread in developmental contexts. Here, we expect tissues to be fragile to the invasion of rapidly dividing and self-renewing mutant cells. This phenomenon has been experimentally demonstrated, for example, after CycD + CDK4 overexpression in neurogenic progenitors [26], and may also be relevant for various mutants in the context of cell competition during development [74].

The model thus predicts that, while the RD topology can respond to perturbations with rapid cell division, it is highly fragile to the invasion of mutant cells (Figure 3C). Mutants with activating mutations (stronger gain) will accumulate due to their strong growth advantage resulting in rapid hyperplasia and clonal dominance compared with the DR topology, where these processes occur much more slowly. There is thus an inherent trade-off between the ability of a topology to reject mutants and its ability to support growth and recovery from large perturbations. This trade-off can be crucial to determine which topology would appear in specific physiological contexts.

## Discussion

The mechanisms that regulate tissue development and homeostasis are complex and involve many signalling, cell identity, and cell division pathways. To make progress in understanding their dynamics, it is important to identify simplifying operating principles. Here, motivated by a range of experimental phenomenology, we developed such an approach by studying the interaction between the cell cycle and cell fate decisions. We propose that regulation of cell fate by G1 lengthening occurs through a saddle-node bifurcation that, through cell-cell interactions, is self-tuned to the vicinity of its critical point. This mechanism provides unique benefits for maintaining a robust set point while leading to the rejection of mutant cells. This mechanism also presents trade-offs that may underpin major physiological changes in development, homeostasis, and ageing.

While the model is based on a well-established molecular mechanism for triggering the G1-S transition (the Rb/E2F bistable switch), its underpinnings are generic and based on universal properties of systems undergoing a saddle-node bifurcation. The effects of mitogens and factors such as cyclins and cell cycle inhibitors are all captured by an effective control parameter that is self-tuned through cell-cell interactions. From the point of view of modelling, this allows us to capture the dynamics of a complex regulatory network by a minimal effective theory. The critical nature of the mechanism also provides the system with a great deal of redundancy, allowing it to integrate many growth signals into a single effective control parameter. Moreover, it allows the circuit to function properly even upon the deletion of specific factors. The mechanism also has significant advantages for signal processing, as it allows for the temporal integration of mitogenic signals over prolonged timescales, thus providing a useful decision-making mechanism in noisy environments [58, 75–79].

The critical lengthening mechanism differs from previously proposed G1 lengthening mechanisms, which rely on time-dependent changes in circuit parameters [44, 80, 81]. In these models, as G1 progresses, circuit parameters gradually shift, eventually causing the system to cross the critical threshold required for the G1-S transition. Both frameworks can be captured by Eq. (2) by introducing a time-dependent distance from bifurcation, *µ*(*t*) (Figure 4). However, the two models exhibit fundamentally different behaviours and make distinct predictions. A strong time-dependence of *µ* would imply weak sensitivity to input signals and to cellular variability, and thus provides an ideal mechanism for synchronous and oscillatory cell divisions [82]. This stands in stark contrast to the properties of the critical lengthening mechanism, which are highly sensitive to cellular variation. Notably, the time-dependent model does not account for experimental observations on the strong sensitivity of G1 length. The experimental observations of Rb protein decreasing in early G1, followed by a relative plateau [83], suggest that both strategies may be employed by cells. Low cell-cell variability may dominate for short G1 lengths due to the time-dependence of the critical point, while critical lengthening and its associated benefits may prevail in slow-cycling cellular populations.

**Figure 4.**
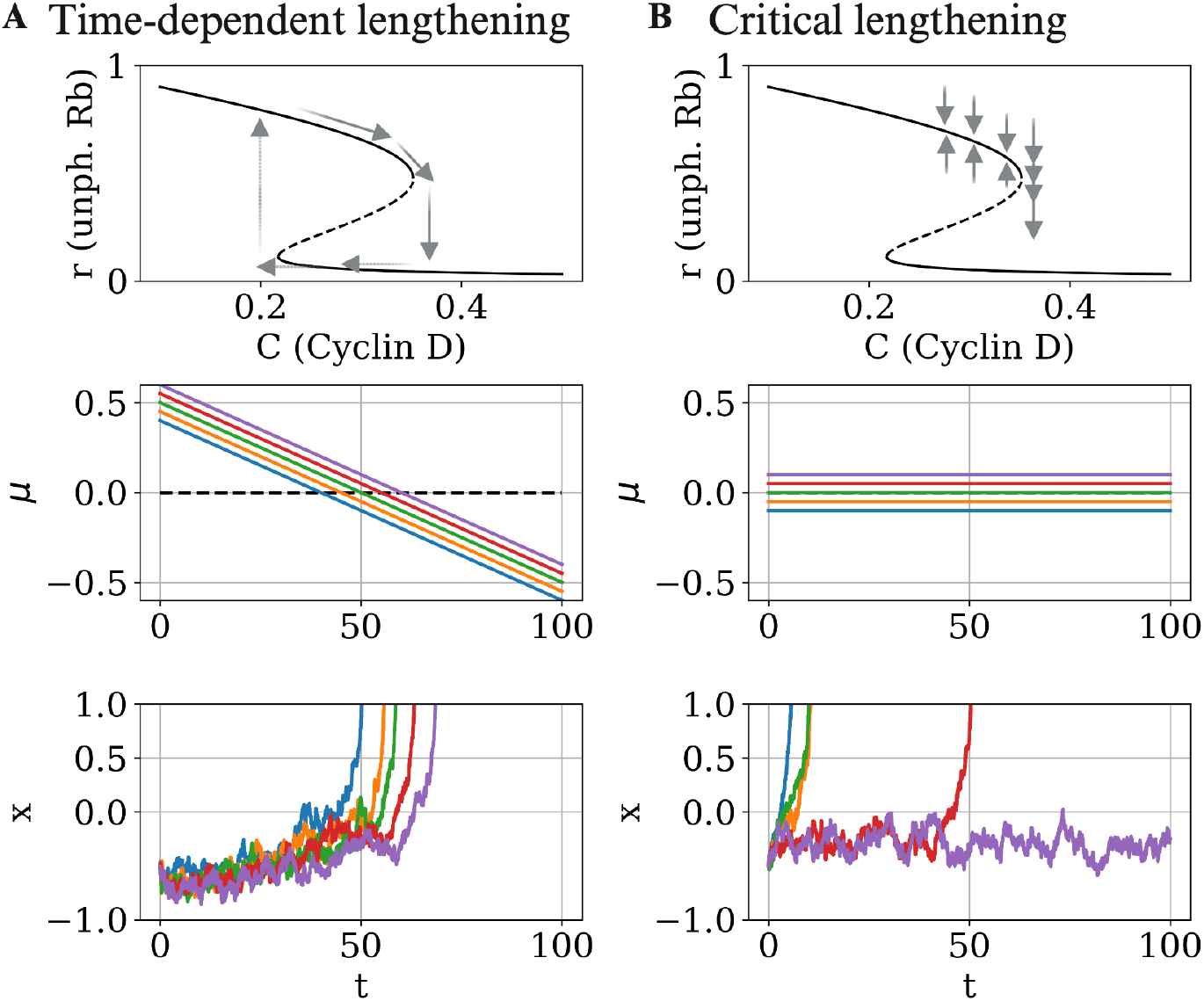
Comparison of critical versus time-dependent G1 lengthening. (A) Time-dependent G1 lengthening is associated with a strong time-dependence of *µ*, the distance from the critical point. (B) By contrast, in the critical lengthening regime, *µ* is held relatively constant over long timescales. The lower panels show a comparison of the two mechanisms, respectively, illustrated by simulations of Eq. (2). The results demonstrate the divergence of G1 lengths associated with the critical lengthening regime, which underlies rapid responses to perturbations and rejection of mis-sensing mutants. The dynamics were simulated with a noise strength of *ξ* = 0.1.

At the population level, the dynamics of the model leads to a self-renewing steady-state associated with finetuning of the underlying molecular rates near the critical point. As such, it relates to previously demonstrated mechanisms of self-tuning to a critical point by competition in the context of transgenerational epigenetic inheritance of gene silencing and immune cell survival [58, 59]. However, unlike previous studies, this mechanism is associated with the steady-state levels of a single self-renewing population, rather than representing a balance between arrival and removal of competing entities. As in earlier work, the key to self-tuning is the sharp sensitivity of the delay around the critical point, which ensures that any extended division time must occur in its vicinity. This sensitivity amplifies variations between cells, allowing for rapid responses and efficient rejection of mutant cells.

The unique sensitivity of G1 length at the bifurcation point allows tissues to have both a very low steadystate turnover and high responsiveness to perturbations. One system where this may be especially crucial is the regulation of beta cell mass in the pancreas. Beta cells have exceptionally low turnover in adults (less than 0.1% per day) [84, 85], but can increase their replication rate rapidly upon the induction of insulin resistance [86], allowing for physiological adaptation [87]. This rapid response can be interpreted using the current model. The primary mitogen for beta cells is glucose, which progresses the G1-S transition by upregulating cyclin D expression [88]. We, therefore, propose the following model for beta cell dynamics: when glucose levels are low, the cell cycle is extended in the stable regime, and the beta cell mass is quiescent - unchanging. An elevation in glucose levels transitions the system to the unstable regime, which is associated with rapid cell divisions, until glucose levels are again lowered, and the system returns to the stable regime. In this system, there is biphasic regulation, where rapidly dividing cells are eliminated, allowing for mutant cell rejection, but conferring risk to glucotoxicity and diabetes [8]. This mechanism is predicted to exploit the sharp sensitivity around the critical point to generate sharp transitions between quiescence and expansion and to remove mis-sensing mutants.

Similar mechanisms may underpin quiescence in adult tissues dominated by the DR topology. Prolonged G1 is a “soft” direction as cells have very slow growth rates. The system may thus transition to this state, only to be reactivated when cues for differentiation transition it back to the unstable phase. The RD topology, on the other hand, does not generate a similar fraction of quiescent cells as, in this case, the tail of the distribution is associated with cells destined for removal. The strong sensitivity of the dynamics around the critical point allows the population to induce quiescence in specific subpopulations, such as DNA-damaged cells, while retaining normal growth for other cells, in line with experimental observations [89].

While our analysis was presented in a relatively simple homeostatic setting, it can be readily extended to other settings, including the regulation of cellular composition in growing cell populations, as occurs, for example, in the pre-implantation embryo [90]. This case is compatible, for example, with feedback from the concentration of one cell type to another. In homeostatic settings, there may also be feedback on the size of the pool of differentiated cells, rather than on the size of the self-renewing compartment. Finally, it is important to note that, while our theory focused on G1 lengthening, its main tenets apply to any regulation based on temporal modulation of transient states, including, for example, the modulation of other cell cycle stages or of states unrelated to the cell cycle.

In addition to modelling biological systems, understanding mechanisms for the regulation of cellular dynamics is crucial for the application of synthetic biology in health and industrial settings, where robust control and mutant cell elimination are central challenges [4, 6, 7, 91, 92]. We propose that, due to its prevalence in natural systems and the clear functional benefits analysed in this study, the regulation of cell fate decisions by temporal modulation may provide an effective design principle for cell population control.

## Methods

### Simulation

The parameters used in the numerical simulations of the competition model are *γ*_*D*_ = 1, *k*_1_ = 1, and *k*_2_ = 0.1, all in arbitrary units. *C* was drawn according to *C* ∼ Norm(*C*_*m*_, *C*_*s*_), with *C*_*s*_ set in proportion to the relative activity of the Wnt pathway. Namely, *C*_*s*_ = 0.05 and *C*_*m*_ = 0.46 (mTeSR1), *C*_*m*_ = 0.57 (E8+IWP2), *C*_*m*_ = 0.65 (E8) and *C*_*m*_ = 1.43 (mTeSR1+Wnt). The G1-S transition was assumed to occur when the level of unphosphorylated Rb dropped below *r <* 0.05. All stochastic simulations were performed using the Euler–Maruyama method.

For cell fate decisions, we assumed, within the RD topology, that the decision to differentiate occurs if G1 length exceeds a number drawn from an exponential distribution with a scale parameter of 20 hours, as plotted in Figure 2. Similarly, in the DR topology, this threshold determined whether a cell undergoes duplication (self-renewal).

### Generalized model for G1 lengthening as a noisy saddle-node bifurcation

In the main text, we considered a minimal model of the G1-S transition based on the concentration of active Rb protein. More generally, we can model the length of G1 using the following general stochastic dynamics,

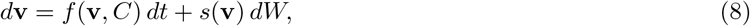

where **v** = *v*_1_ … *v*_*K*_ represents the state of a gene regulatory network (GRN), *f* = *f*_1_ …, *f*_*K*_ defines the dynamics of the GRN, *s* = *s*_1_, …, *s*_*K*_ defines the state-dependent amplitude of the noise, *W* is an *K*-dimensional Wiener process, and *C* is a control parameter for a saddle-node bifurcation that destabilizes the G1 state, denoted **v**_*G*1_. While *f* can correspond to the dynamics of the Rb/E2F/CycE network and *C* to the concentration of CycD, the general analysis does not make specific assumptions about the number of components or dynamics of the GRN, allowing us to make general conclusions that extend to other possible molecular mechanisms. In this sense, Eq. (1) is a specific example of Eq. (8).

In an experimental setting, there may be slight differences in biochemical parameters or sensed environments between individual cells. In our model, we can capture this by indexing the GRN of each cell as **v**_*i*_ (with *i* = 1, 2, …) and setting *C*_*i*_ as a random variable. The dynamics of a cell population are then given by

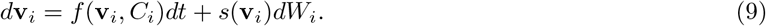

Variation in the control parameters *C*_*i*_, as well as noise, cause cell-to-cell differences in the length of G1. We can also think of variation in *C*_*i*_ as fluctuations on the timescale of cell growth and division, while *W* captures fluctuations on a faster timescale. In the vicinity of the critical point, the dynamics of Eq. (9) are captured by the stochastic normal form, Eq. (2), where the GRN coordinates and time are rescaled to *x* and 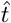, while *ξ* corresponds to the fluctuations near the critical point, and *µ*_*i*_ is the (rescaled) distance from bifurcation of cell *i*. More formally, *µ*_*i*_ ∝ *C*_*i*_ − *C*_crit_. The duration of G1 is then given by [57]

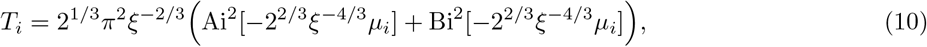

where Ai, Bi are Airy functions of the first and second kind, respectively. There is a steep dependence of *T* on the bifurcation parameters, so that *T*_*i*_ increases super-exponentially as *µ*_*i*_ decreases (and thus as *C*_*i*_ drops below the critical value).

### Population dynamics

Consider a population of cells with identical cell-cycle duration *T*, and let *P*_*R*_(*T*) define the probability that a cell division after duration *T* concludes in cell duplication (self-renewal), while *P*_*D*_(*T*) defines the probability that it concludes in differentiation; we assume that cell fate decisions are symmetric and thus *P*_*D*_(*T*) = 1 − *P*_*R*_(*T*). Thus, after each time interval *T*, the population doubles, contributing a net growth of (*P*_*R*_(*T*) − *P*_*D*_(*T*))*N* = (2*P*_*R*_(*T*) − 1)*N* cells to the population. Thus, defining

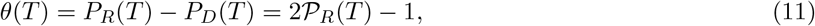

we have that the population dynamics, for cells with an (identical) cycle length *T*, is given by:

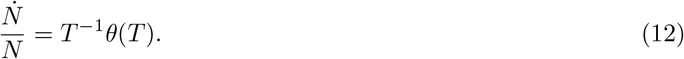

## Acknowledgments

We wish to thank Alexis Barr for helpful discussions and critical reading of the manuscript. B.D.S. acknowledges the support of the Royal Society through an EP Abraham Research Professorship (RSRP/R/231004). For the purpose of open access, the authors have applied a Creative Commons Attribution (CC BY) licence to any Author Accepted Manuscript version arising from this submission.

## Author contributions

The project was conceived, developed, and the manuscript prepared by O.K. with continuous input from B.D.S.

